# Motor training strengthens corticospinal suppression during movement preparation

**DOI:** 10.1101/2020.02.14.948877

**Authors:** Pierre Vassiliadis, Gerard Derosiere, Julien Grandjean, Julie Duque

## Abstract

Training can improve motor skills and modify neural activity at rest and during movement execution. Learning-related modulations may also concern motor preparation but the neural correlates and the potential behavioral relevance of such adjustments remain unclear. In humans, preparatory processes have been largely investigated using transcranial magnetic stimulation (TMS) with several studies reporting decreased corticospinal excitability (CSE) relative to a baseline measure; a phenomenon called preparatory suppression. Here, we investigated the effect of motor training on preparatory suppression of CSE in humans, as compared to modulatory changes at rest. We trained participants to initiate quick movements in an instructed-delay reaction time (RT) task and used TMS to investigate changes in CSE over the practice blocks. Training on the task speeded up RTs, with no repercussion on error rates. Training also increased baseline CSE at rest. Most interestingly, we found that motor activity during action preparation did not mirror the training-related rise in resting CSE. Rather, the degree of preparatory suppression from the rising baseline strengthened with practice. This training-related change in preparatory suppression predicted RT gains (but not the changes in baseline CSE): subjects showing a stronger expansion of preparatory suppression were also those exhibiting larger gains in RTs. Finally, such relationship between RTs and preparatory suppression was also evident at the single-trial level: RTs were generally faster in trials where preparatory suppression was deeper. These findings suggest that training induces changes in motor preparatory processes that are linked to an enhanced ability to initiate fast movements.

**New and Noteworthy:** Any movement is preceded by a period of preparation, which involves a broad suppression of the corticospinal pathway, a phenomenon called preparatory suppression. Here, we show that motor training strengthens preparatory suppression and that this strengthening is associated with an acceleration of movement initiation. Our findings yield an extension of former work, highlighting a key role of preparatory suppression in training-driven behavioral improvements.

## 1. Introduction

Motor training improves the speed and/or accuracy at which movements are selected, initiated and executed. Significant research has been devoted to unveiling the functional changes at the basis of such improvements (Krakauer et al. 2019). At the neural level, neuroimaging (*e.g.*, Wiestler & Diedrichsen, 2013; Wenger et al., 2017; Yokoi & Diedrichsen, 2019) and transcranial magnetic stimulation (TMS) studies (*e.g.*, Rosenkranz et al., 2007; Reis et al., 2008; Mawase et al., 2017) have shown that training is accompanied by a plastic reorganization of the motor system, supporting the formation of new motor memories. Specifically, training amplifies resting motor activity (*e.g.*, Pascual-Leone et al., 1995; Butefisch et al., 2000; Duque et al., 2008; Galea & Celnik, 2009; Christiansen et al., 2018) and induces learning-specific changes of motor activity during movement execution (Krakauer et al. 2004; Shmuelof et al. 2014; Steele and Penhune 2010). Animal studies also show learning-related modulations of motor activity during action preparation (Makino et al. 2017; Paz et al. 2003; Vyas et al. 2018, 2020) that could reflect an optimization of preparatory processes with training (Mawase et al. 2018). Yet, the behavioral relevance of the effects of training on action preparation remain unclear.

In humans, the excitability of the motor system can be assessed by applying TMS over primary motor cortex (M1), eliciting motor-evoked potentials (MEPs), whose amplitude reflects the excitability of the corticospinal pathway (Derosiere et al. 2020; Derosiere and Duque 2020). When applied during reaction time (RT) tasks, TMS elicits MEPs that are used to assess corticospinal excitability (CSE) changes associated with action preparation and initiation. CSE is often suppressed during action preparation when compared to a baseline, measured at rest. The function of this preparatory suppression (or inhibition) remains unclear (*e.g.*, Greenhouse et al., 2015; Duque et al., 2017; Derosiere, 2018; Hannah et al., 2018). A prominent view is that it assists action selection processes, by preventing the release of premature or incorrect responses (Duque et al. 2010; Quoilin et al. 2018). Indeed, the amount of preparatory suppression was shown to scale with the complexity of selection processes (Duque et al. 2016; Klein et al. 2014). Another hypothesis is that preparatory suppression eases action initiation (Greenhouse et al. 2015; Hasegawa et al. 2017). In this line, a study showed a dependence of RTs on the amount of preparatory suppression on a single-trial basis: the stronger the suppression, the faster the initiation of the ensuing movement (Hannah et al. 2018). Importantly, these hypotheses could be both valid as they focus on different levels of control, which are both known to shape motor activity: while the choice hypothesis suggests that suppression originates from processes that help select accurate actions (*i.e.*, therefore reducing the error rate), the motor hypothesis entails that preparatory suppression is also generated by processes speeding up action initiation (*i.e.*, therefore reducing RTs).

Here, we investigated the impact of motor training on preparatory suppression, while subjects practiced an instructed-delay RT task. The choice aspects were clear-cut, as evident from the low error rates, even before training. Hence, in such task, the selection requirements are so low that there is no room for improvement; subjects can only become more skilled at the motor level, by initiating their action faster. Based on this, we predicted that RTs would decrease over the course of practice but that error rates would remain marginal. In addition, we expected resting CSE to increase with training, in accordance with previous work (Butefisch et al. 2000; Christiansen et al. 2018; Duque et al. 2008; Galea and Celnik 2009; Pascual-Leone et al. 1995). Based on the motor hypothesis (*i.e.*, that preparatory suppression fastens RTs), we expected that training would deepen the drop in excitability during action preparation with respect to rest, reflecting an increased preparatory suppression. Hence, we predicted that preparatory activity would not follow the training-related rise in resting CSE.

## 2. Materials and Methods

### 2.1. Participants

Fifteen right-handed healthy subjects participated in the present study (n=15; 10 women; 22.4±1.63 years old). Handedness was assessed via Edinburgh Handedness inventory (Oldfield 1971). Participants filled out a TMS safety questionnaire to look for any contra-indication and gave written informed consent in accordance with the Ethics Committee of the Université Catholique de Louvain and the principles of the Declaration of Helsinki. We had to exclude one subject because we encountered a technical problem during the experiment; hence, analyses were run on the fourteen remaining subjects. Part of the data reported here has been exploited in a separate study (Vassiliadis et al. 2018). All of the data are expressed as mean±SE.

### 2.2. Task

Subjects were sited in front of a computer screen with the hands on response devices **(Fig.1A**, (Quoilin et al., 2016, 2018, 2019; Grandjean et al., 2019). They performed an instructed-delay RT task, which required them to choose between abduction movements of the left or right index finger according to the position of a preparatory cue (*i.e.*, a left- or right-sided ball separated from a goal by a gap). Participants had to prepare their movement once the ball appeared but to withhold responding until the onset of a “Go” signal (*i.e.*, a bridge). When the bridge appeared on the screen, subjects had to respond as fast as possible, allowing the ball to roll on the bridge and to reach the goal. To reduce anticipation of the “Go” signal, the bridge did not appear in some of the trials (5%). Trials always ended with a feedback score reflecting performance (see **Fig.1B**).

**Figure 1.**
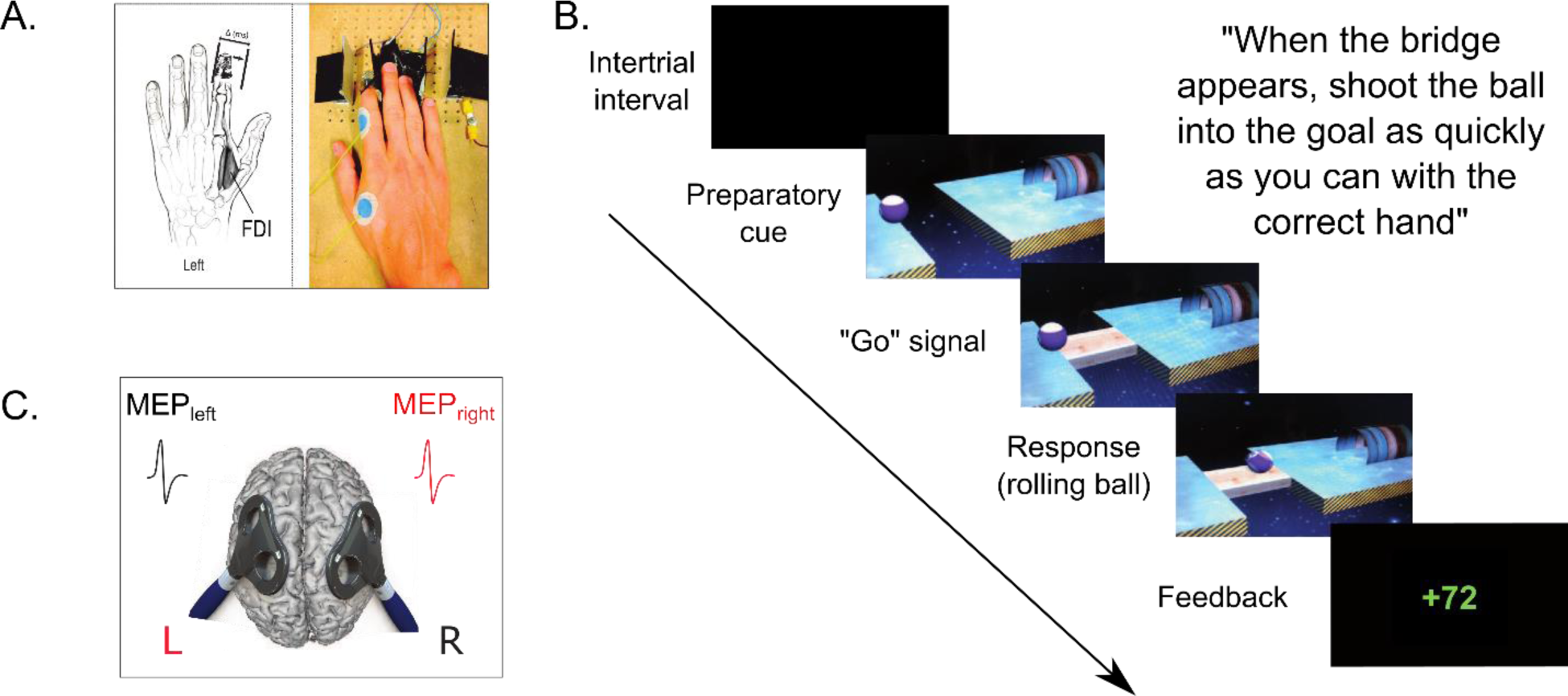
A, The response device. Index finger responses were recorded using a home-made device composed of two pairs of metal edges fixed on a wooden platform and positioned under the left (graphic representation) and right (photographic representation) hands. **B, “Rolling Ball” task.** Subjects were asked to choose between responding with the left or right index finger according to the position of a ball (Preparatory cue) appearing on the left or right part of the screen (left in the current example). They had to wait until the onset of a bridge (Go signal) to release their response as quickly as possible. The ball then rolled on the bridge (when the subjects answered correctly) to reach a goal located on the other side of the gap. A feedback reflecting how fast and accurate the subjects were concluded each trial. On correct trials, scores ranged from 1 to 100 points and were displayed in green. Participants were informed that the score was inversely proportional to the RT: the faster the response, the higher the score. In order to homogenize the score across subjects, scores on correct trials were individualized according to RTs measured during a familiarization session just before the main experiment (Vassiliadis et al., 2018; Grandjean et al., 2019). Incorrect responses were penalized with negative scores displayed in red. They involved responses occurring too early (RT<100 ms), referred to as “anticipation errors” (−75 points), responses occurring too late (RT>500 ms), referred to as “time-out errors” (−50 points), responses provided with the incorrect hand (−20 points), referred to as “choice errors” and responses provided on catch trials (−12 points), referred as “catch errors”. When subjects succeeded not to respond on a catch trial, they were rewarded by +12 points. The total score was displayed at the end of each block. **C, TMS protocol.** Two figure-eight-shaped coils were placed over the subject’s M1, eliciting MEPs in the left and/or right FDI. In double-coil trials, a 1-ms interval separated the onset of the two pulses, eliciting MEPs in both hands at a near simultaneous time (Algoet et al. 2018; Grandjean et al. 2018; Quoilin et al. 2019; Vassiliadis et al. 2018). This interval was used to avoid direct electromagnetic interference between the two coils (Cincotta et al. 2005), while preventing transcallosal interactions that would occur between motor areas with longer delays (Ferbert et al. 1992; Hanajima et al. 2001). Notably, in double-coil trials, half of the trials involved a pulse over left M1 first whereas the other half involved a pulse over right M1 first (1ms delay). These data were assembled because the a prior analysis reported elsewhere showed that the order of stimulation does not influence the double-coil MEP amplitudes, which were identical to single-coil MEPs (Vassiliadis et al. 2018).

### 2.3. TMS Protocol

Monophasic pulses were delivered through one or two figure-of-eight shaped coils, each connected either to a Magstim 200^2^ magnetic stimulator (Magstim, Whitland, Dyfed, UK) or a Magstim Bistim^2^ magnetic stimulator. The TMS machine used to stimulate each hemisphere was counterbalanced between subjects. Pulses could be triggered in one (*i.e.*, single-coil TMS) or in the two coils (*i.e.*, double-coil TMS, **Fig.1C**) because the dataset was initially acquired for a separate study to establish the reliability of double-coil TMS to probe CSE bilaterally (Grandjean et al. 2018; Vassiliadis et al. 2018). Here, MEPs are considered regardless of the protocol used to elicit them.

Each TMS coil was placed tangentially over one M1 with the handle pointing backward and laterally at a 45° angle away from the midline **(Fig.1C)**. TMS was applied over the hotspot of the first dorsal interosseous muscle (FDI), which was the prime-mover in our task (Duque et al., 2014; Derosiere et al., 2017a, 2017b). The resting Motor Threshold (rMT) was determined for each M1. It was defined as the minimal intensity required to evoke MEPs of 50µV at rest in at least 5 out of 10 stimulations. The rMTs equalled 41.7±5.05% and 40.8±6.39% of the maximum stimulator output for the left and the right FDI, respectively. For each hemisphere, the intensity used throughout the experiment was set at 115% of the individual rMT (Derosiere et al. 2019).

### 2.4 Experimental procedure

The experiment started with two familiarization blocks. The first block allowed subjects to become acquainted with the task. The second block involved TMS and served to compute the median RT for each subject. The latter was used to individualize the feedback scores on correct trials according to the initial performance (see **Fig.1B**).

Then, subjects performed 400 trials of the task, divided in 10 blocks. Each block involved an equal combination of single- and double-coil stimulations, occurring in a random order (*i.e.*, subjects could not anticipate the type of stimulation they would face). Given that both techniques produce equivalent MEPs (Grandjean et al. 2018; Vassiliadis et al. 2018), these data were considered regardless of the protocol used to elicit them.

TMS could occur in three different settings. First, some TMS pulses were delivered outside the blocks (TMS_baseline-out_), providing MEPs reflecting baseline CSE at complete rest. TMS_baseline-out_ pulses occurred every other block, starting before block 1 and ending after block 8 (5 time points; 30 MEPs per time point; **Fig.2A**). Second, TMS occurred during the intertrial interval, 300 ms before the beginning of the trial (**Fig.2B**). MEPs recorded at this time (12 per block) provided another baseline measure of CSE, with subjects at rest but engaged in the task (TMS_baseline-in_ Labruna et al., 2011). Finally, TMS occurred at 900 or 950 ms after the onset of the preparatory cue (TMS_preparation_). Since no difference was found between MEPs recorded at these two timings in our previous analysis (Vassiliadis et al. 2018), these data were pooled together (48 MEPs per block). Half of these MEPs fell in left response trials, while the other half occurred in right response trials. Hence, MEPs could either fall in a hand that was selected for the forthcoming response (MEP_selected_; *e.g.*, left MEPs preceding a left index finger response) or in a hand that was non-selected (MEP_non-selected_).

**Figure 2.**
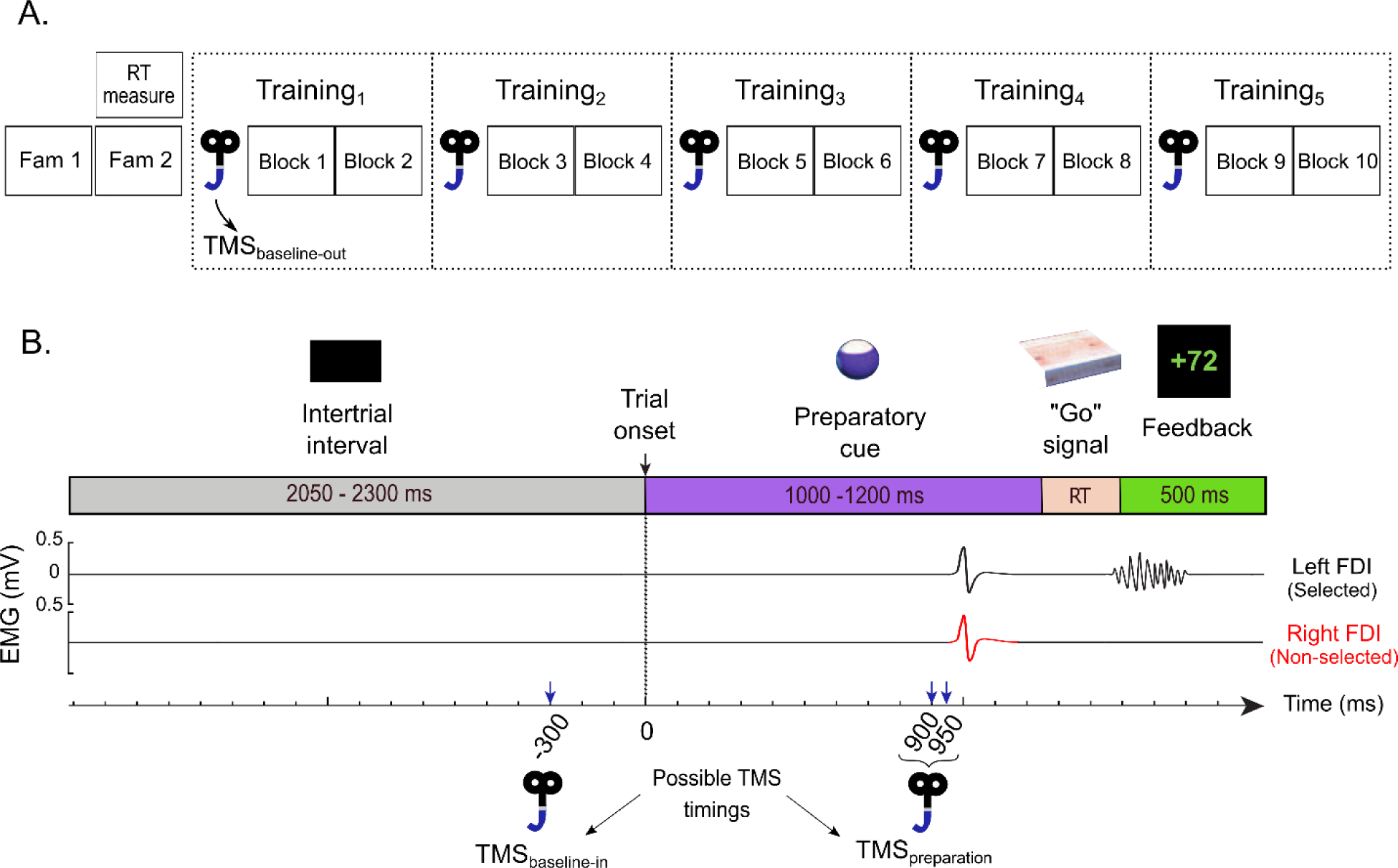
A, Time-course of the experiment. After two familiarization blocks (Fam1: 20 trials without TMS and Fam 2: 40 trials with TMS), subjects executed ten blocks of forty trials during which MEPs were elicited at TMS_baseline-in_ or TMS_preparation_ (see 2A). The effect of training was assessed by comparing five sets of data (Training_1_ to Training_5_), each involving MEPs pooled over two consecutive blocks. MEPs were also elicited outside the blocks (TMS_baseline-out_) at five points in time, before block 1 and after blocks 2, 4, 6 and 8, categorized as Training_1_ to Training_5_, similar to the MEPs elicited during the blocks. We obtained 15 TMS_baseline-out_ MEPs per hand for each Training_STAGE._ Comparing these data sets allowed us to consider potential training-related changes in resting CSE outside the context of the task. **B, Time course of a trial.** Trials were separated by a blank screen (intertrial interval; 2050 to 2300 ms) and always started with a preparatory cue appearing for a variable delay period (1000 to 1200 ms). Then, a Go signal was presented and remained on the screen until a response was detected, hence for the duration of the reaction time (RT). The feedback was presented at the end of each trial for 500 ms and depended on the RT on correct trials. Importantly, the variable delays in the task were sampled from uniform distributions to induce temporal uncertainty and therefore reduce anticipation that could emerge with the repetition of trials. TMS pulses occurred either during the intertrial interval (300 ms before the beginning of the trial; TMS_baseline-in_), or during the delay period (900 or 950 ms after the preparatory cue onset, timings pooled; TMS_preparation_). In double-coil trials, motor-evoked potentials (MEPs) were elicited in the first dorsal interosseous (FDI) of both hands at a near simultaneous time (1 ms delay); in single-coil trials, MEPs were elicited in the left or right hand. The figure displays a left hand trial with double-coil TMS at TMS_preparation_.

### 2.5 Data processing and statistical analyses

The purpose of the study was twofold: (1) to characterize changes in CSE at rest and during action preparation occurring along with training in a basic instructed-delay RT task, (2) to assess whether modulations in CSE were correlated to training-related improvements in RTs. To do so, the behavioral and MEP data were evaluated according to the block within which they were elicited and data from two consecutive blocks were pooled together. Given the 10 blocks, we obtained five data sets reflecting five training stages (Training_STAGE_: Training_1_ to Training_5_; **Fig.2A)**.

Statistical analyses were carried out with Matlab 2018a (the Mathworks, Natick, Massachusetts, USA) and Statistica 10 (StatSoft Inc., Tulsa, Oklahoma, USA). All data were systematically tested for the sphericity assumption using Maunchley’s tests. The Greenhouse– Geisser (GG) correction was used for sphericity when necessary. Post-hocs comparisons were always conducted using the Fisher’s LSD procedure. The significance level was set at p ≤ 0.05.

#### 2.5.1 RTs and errors

Left and right hand RTs were computed as the difference between the onset of the Go signal and movement onset (when the finger quitted the outer metal edge of the device). Trials where subjects made an error were removed from the data set for the RT analysis. An average of 35 left and 34 right response trials remained for each subject at each Training_STAGE_. We computed the mean RT for left and right responses separately and then averaged these data together. Besides, we also assessed response accuracy over training, by computing, for each Training_STAGE_, the amount of anticipation, time-out and catch errors as well as the total error rate. For each of these variables, we expressed the number of incorrect trials in percentage of the total amount of trials, regardless of the responding hand. Choice errors were not analysed because they were rare (4 choice errors across all subjects). For the statistical analysis of RTs and errors (*i.e.*, anticipation, time-out, catch and global errors), we used one-way analyses of variance for repeated measures (ANOVA_RM_) with the factor Training_STAGE_ (Training_1_ to Training_5_).

#### 2.5.2 MEP amplitudes

MEPs were obtained by recording electromyography (EMG) bilaterally from surface electrodes (Neuroline, Medicotest, Oelstykke, Denmark) placed over the FDI. The signals were amplified (x1000), bandpass filtered (10-500Hz; NeuroloLog; Digitimer), digitalized at 2000 Hz and collected with Signal (Signal 3.0, Cambridge, UK) for offline analysis. Trials with background EMG activity preceding the pulse exceeding 3 SDs above the mean were discarded (1.68±0.30% removal; Vassiliadis et al., 2018; Grandjean et al., 2018, 2019). This was done to prevent contamination of the MEP measurements by significant fluctuations in background EMG.

To assess training-related changes in resting CSE based on MEPs elicited at TMS_baseline-out_ and TMS_baseline-in_, we averaged separately left and right hand MEPs for each Training_STAGE_ before computing the mean of these averages. These data were analysed using a two-way ANOVA_RM_ with TMS_TIMING_ (TMS_baseline-out_ or TMS_baseline-in_), and Training_STAGE_ (Training_1_ to Training_5_) as within-subject factors. To assess training-related changes in preparatory suppression based on MEPs at TMS_preparation_ (expressed in percentage of MEPs at TMS_baseline-in_), we first removed the trials in which subjects made a mistake (10.78±1.81% removal) and then grouped left and right hand MEPs according to whether they corresponded to a MEP_selected_ or MEP_non-selected_. Within these categories, we averaged the separate means of left and right hand MEPs for each Training_STAGE_. To analyse these data, we first focused on percentage MEPs at Training_1_, assessing with t-tests (against a constant value of 100%) the significance of preparatory suppression at the beginning of training. Then, we analyzed all training stages using a two-way ANOVA_RM_ with the factors MEP_SELECTION_ (MEP_selected_ or MEP_non-selected_) and Training_STAGE_ (Training_1,_ to Training_5_). This ANOVA was also run on absolute MEP amplitudes (in mV).

#### 2.5.3 Relationship between training-related changes in RTs and CSE

As described in the Result section, training influenced RTs and CSE. We studied the relationship between changes at these two levels, with CSE considered separately at rest and during action preparation. We computed ratios reflecting training-related changes. Based on the RT data, we realized that the subjects’ behavior improved substantially during the first practice stage (Training_1_ to Training_3_) but then, RTs remained quite stable (from Training_3_ to Training_5_; Result section). For this reason, we considered ratios for these two phases of training separately, providing us with an indication of early (Training_ratio-early_: Training_3_/Training_1_) and late (Training_ratio-late_: Training_5_/Training_3_) training-related changes in RTs and CSE. For the latter, we computed separate ratios for MEPs at TMS_baseline-out_, TMS_baseline-in_ and TMS_preparation_ (expressed in percentage of MEPs at TMS_baseline-in_). We then examined the correlation between the RT and MEP Training_ratios_ by using least squares linear regressions.

Finally, we compared the strength of the RT relationship to training-related changes in MEP amplitudes at TMS_baseline-in_ (reflecting resting CSE) and changes in percentage MEPs at TMS_preparation_ (reflecting preparatory suppression of CSE). To do so, in order to obtain a robust estimate of the absolute Pearson’s R, we ran a bootstrap analysis with 10000 resamples and calculated a median R for each correlation (Efron 1979). These R-values were then compared to each other using Pearson and Fillon’s z test (Pearson and Filon 1898).

#### 2.5.4 Single-trial relationship between RTs and preparatory suppression

The correlation analyses revealed a relationship between RTs and preparatory suppression: the subjects who showed the greatest training-related reduction in RTs were also those who displayed the strongest deepening in preparatory suppression (see Result section). To better understand the dependency of RTs to the strength of preparatory suppression, we investigated whether this relationship was evident on a single-trial basis, as suggested previously (Hannah et al. 2018). We selected the MEPs elicited at TMS_preparation_ and again, expressed them as a percentage of TMS_baseline-in_. We only used the double-coil trials, to consider a homogeneous set of data, with preparatory MEPs falling in both hands systematically. For each trial, we extracted the RT, as well as the MEPs recorded at TMS_preparation_ in both selected (MEP_selected_) and non-selected (MEP_non-selected_) hands. Hence, for each trial, we obtained one RT measure linked to two different MEPs.

To examine the relationship between RTs and preparatory suppression, we pooled the trials from all 10 blocks together and sorted them according to the amplitude of MEPs within each trial. Given that there were two MEPs in each trial, we repeated this procedure twice, providing us with two different orderings of the trials according to the MEP_selected_ or MEP_non-selected_. Within each arrangement, trials were grouped into 6 consecutive percentile bins (MEP_BIN_: MEP_BIN-1_ = 0 to 16.7%, MEP_BIN-2_ = 16.7 to 33.3% …MEP_BIN-6_ = 83.3 to 100% of the data). MEP_BIN-1_ contained the trials with the stronger preparatory suppression whereas the MEP_BIN-6_ included the trials with the weaker preparatory suppression. We then computed the mean RT of trials within each MEP_BIN_ (23 trials per condition on average and never less than 19 trials), and then averaged the corresponding RTs. Hence, we obtained six average RT values (*i.e.*, one for each MEP_BIN_) for each of the trial arrangements based on the two MEP types. These two sets of RT data were analysed using two separate ANOVA_RM_ with the factor MEP_BIN_ (MEP_BIN-1_ to MEP_BIN-6_).

## 3. Results

### 3.1. RTs and errors

**Fig.3A** shows the evolution of RTs with training. The ANOVA_RM_ revealed a significant influence of Training_STAGE_ on RT (F_(4,52)_=4.31, p=0.0043). Post-hoc tests showed that RTs measured from Training_3_ to Training_5_ were shorter than at Training_1_ (all p<0.004). In contrast, the total error rate remained stable over the blocks (F_(4,52)_=0.82, p=0.52, **Fig.3B**). We did not observe any modification of the percentage of anticipation (F_(4,52)_=1.12, p=0.36), time-out (GG-corrected F_(2.50,32.50)_=0.90, p=0.44) or catch errors (F_(4,52)_=1.73, p=0.16). Hence, training enabled subjects to respond more quickly while maintaining the same accuracy level.

**Figure 3.**
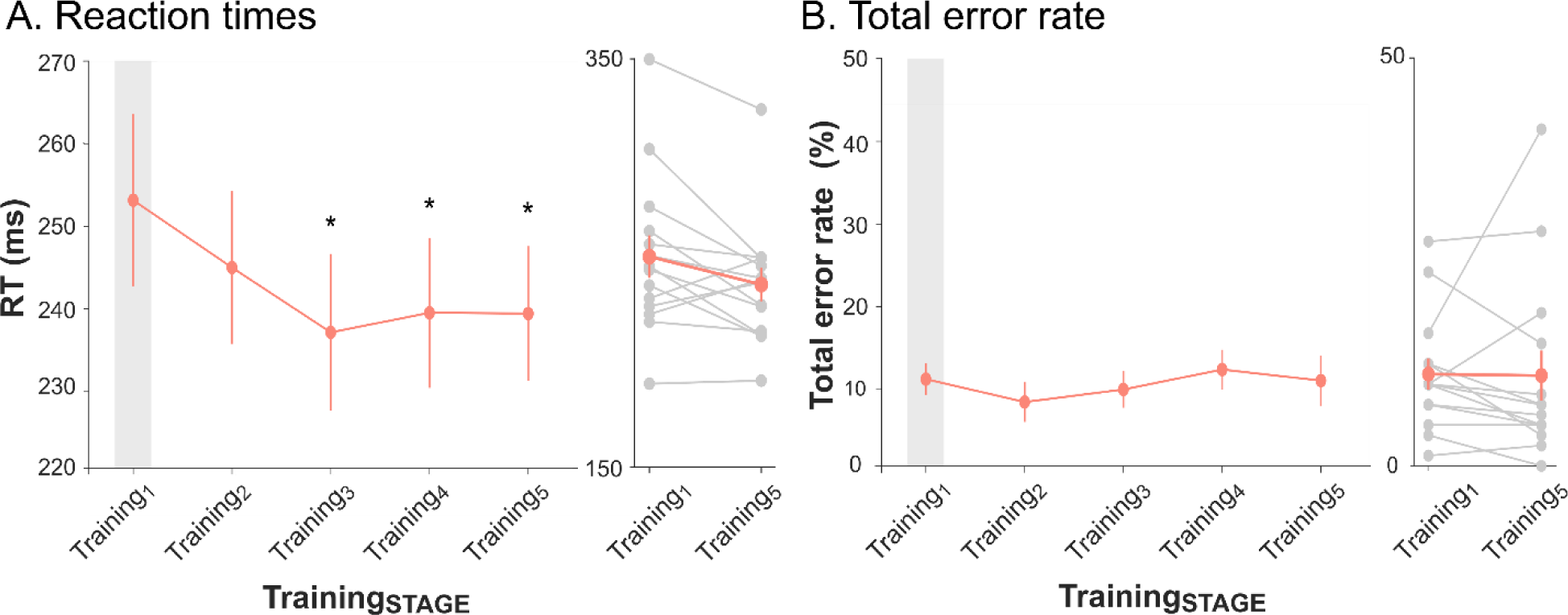
Evolution of reaction times (RTs) and total error rate throughout training. The mean RTs (**A**, in ms) and total error rate (**B**, in % of all trials) are represented for each Training_STAGE_, regardless of the responding hand. Stars denote a significant difference between a given Training_STAGE_ and Training_1_ (p<0.05). Individual data for Training_1_ and Training_5_.

### 3.2. MEP amplitude

First, we evaluated the effect of training on MEPs acquired at rest. As evident on **Fig.4**, MEPs were larger when assessed in the context of the task (TMS_baseline-in_: 1.79±0.17mV) compared to when subjects were fully at rest (TMS_baseline-out_: 1.34±0.17mV), as supported by the significant factor TMS_TIMING_ (F_(1,13)_=28.43, p<0.001) and consistent with previous studies (Derosière et al. 2015; Labruna et al. 2011). The ANOVA_RM_ also revealed an effect of Training_STAGE_ on baseline MEPs (F_(4,52)_=6.34, p<0.001). MEPs recorded at Training_2_ to Training_5_ were larger than at Training_1_ (all p<0.03). This training effect on MEPs occurred independently of the TMS_TIMING_: there was a parallel increase in the amplitude of MEPs elicited at TMS_baseline-out_ and TMS_baseline-in_ (Training_STAGE_xTMS_TIMING_: F_(4,52)_=0.18, p=0.95).

**Figure 4.**
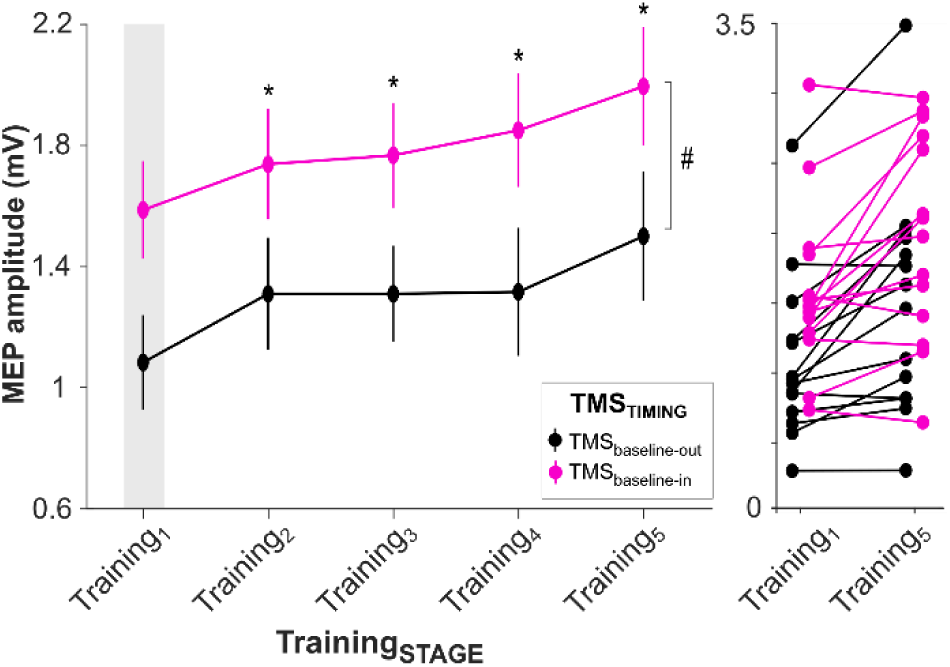
Evolution of baseline MEPs throughout training. MEP amplitudes (in mV) elicited at TMS_baseline-out_ (black) and TMS_baseline-in_ (pink) at the different Training_STAGES_. Hash signs indicate a TMS_TIMING_ effect. Stars denote a significant difference between a given Training_STAGE_ and Training_1_ (p<0.05). Individual data for Training_1_ and Training_5_ are also displayed.

Second, we analyzed the effect of training on preparatory suppression by considering MEPs elicited at TMS_preparation_ (expressed in percentage of TMS_baseline-in_). As evident on **Fig.5A**, percentage FDI MEPs were initially suppressed at Training_1_ (MEPs smaller than 100%), consistent with the presence of preparatory suppression in the prime-mover, whether selected for the forthcoming response (MEP_selected_: 73.98±4.00%; t_(13)_=-6.50, p<0.0001) or not (MEP_non-selected_: 76.26±4.36%; t_(13)_=-5.44, p<0.001). Interestingly, preparatory suppression became more prominent with training: the ANOVA_RM_ revealed a significant decrease in percentage MEP amplitudes over the Training_STAGES_ (F_(4,52)_=2.79, p=0.036). This change was marginal at Training_4_ (*i.e.*, Training_4_: p=0.058 when compared to Training_1_) and became significant at Training_5_ (p=0.006). It concerned MEPs obtained from the selected and non-selected hands (Training_STAGE_xMEP_SELECTION_: F_(4,52)_=0.56, p=0.70). To further our understanding of training-related changes of preparatory activity, we ran another set of ANOVA_RM_ on absolute MEP amplitudes (rather than percentages) at TMS_preparation_ (**Fig.5B**). These MEPs did not show any fluctuation over the Training_STAGES_ (F_(4,52)_=1.30, p=0.28). Moreover, we did not find any MEP_SELECTION_ effect (F_(1,13)_=1.69, p=0.22) or Training_STAGE_xMEP_SELECTION_ interaction (F_(4,52)_=0.85, p=0.50).

**Figure 5.**
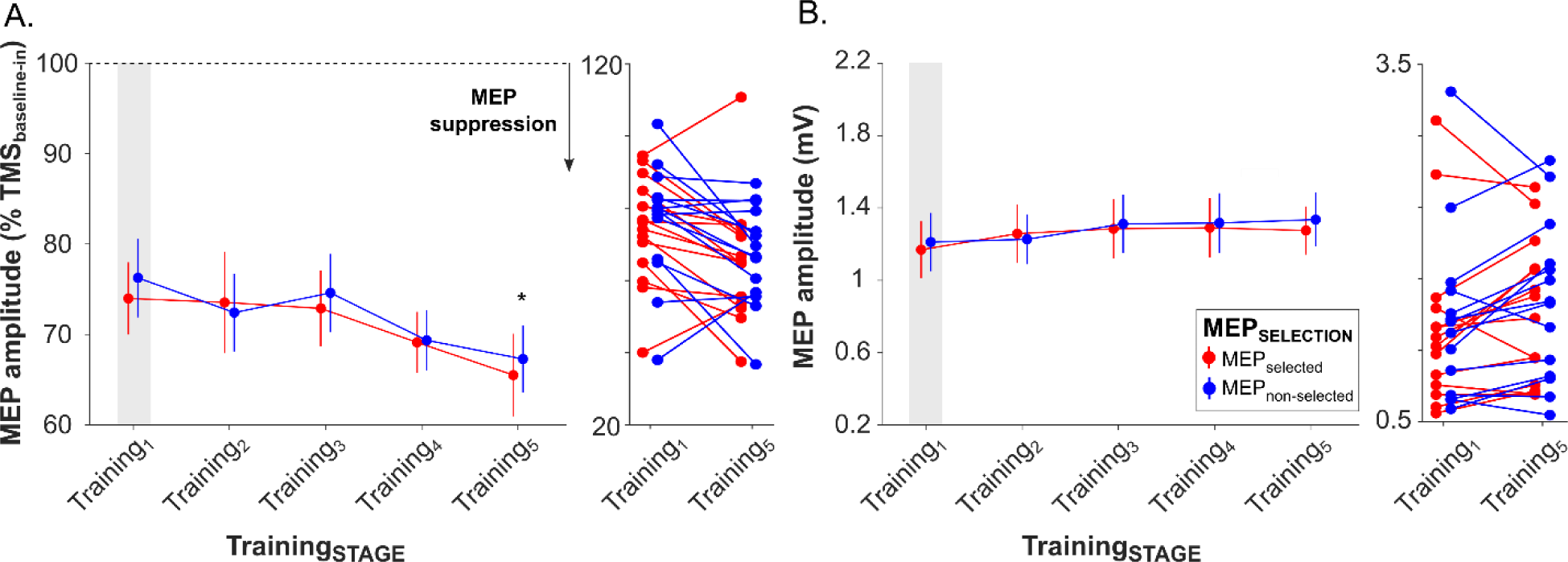
Evolution of preparatory MEPs throughout training. Normalized MEP amplitudes recorded at TMS_preparation_ (in percentage of MEPs elicited at TMS_baseline-in_) muscles at the different Training_STAGES_ **(A)**. Absolute MEP data (in mV) are also represented muscles **(B)**. The star denotes a significant difference between a Training_5_ and Training_1_ (p<0.05). Note that the change in preparatory suppression was close to significance at Training_4_ (*i.e.*, p=0.058 when compared to Training_1_). Individual data for Training_1_ and Training_5_ are also displayed.

In conclusion, our results indicate that training did not produce even modulatory changes in motor activity at rest and during action preparation: while resting CSE increased, preparatory activity remained flat over the blocks, thus revealing an augmenting drop (*i.e.*, preparatory suppression) with respect to the rising baseline excitability state. These changes in CSE occurred in parallel with an acceleration of RTs.

Because RTs became shorter over the blocks, one may argue that MEPs at TMS_preparation_ were not recorded in a comparable preparatory state throughout training; that is, the delay between TMS and movement onset (Delay_TMS-TO-MOVE_) may have decreased over the blocks. Importantly, we shuffled the delay between the pulse and the Go signal in the present study (see Methods), in order to prevent changes in RT to convert into equivalent changes in the Delay_TMS-TO-MOVE_. However, because TMS fell on average closer to movement onset at Training_5_ (399.70±8.48ms) than Training_1_ (419.99±9.99ms, t_(13)_=-3.10, p=0.008), we performed an additional analysis to control for a potential bias of the Delay_TMS-TO-MOVE_. We conducted a response-locked analysis whereby we classified MEP data at TMS_preparation_ (regardless of the Training_STAGE_) according to the Delay_TMS-TO-MOVE_ in 5 consecutive bins of trials (Delay_BIN_ = Delay_BIN-1_= 0 to 20%, Delay_BIN-2_= 20 to 40%, …, Delay_BIN-5_= 80 to 100% of the Delay_TMS-TO-MOVE_ data). An ANOVA_RM_ ran on these data did not reveal any effect of Delay_BIN_ (F_(4,52)_=1.45; p=0.23), nor was there any significant MEP_SELECTION_xDelay_BIN_ interaction (F_(4,52)_=0.40; p=0.81; **Fig.6**). These results indicate that MEPs elicited preceding a Go signal remain quite unaffected by the delay separating the TMS_preparation_ pulse and movement onset.

**Figure 6.**
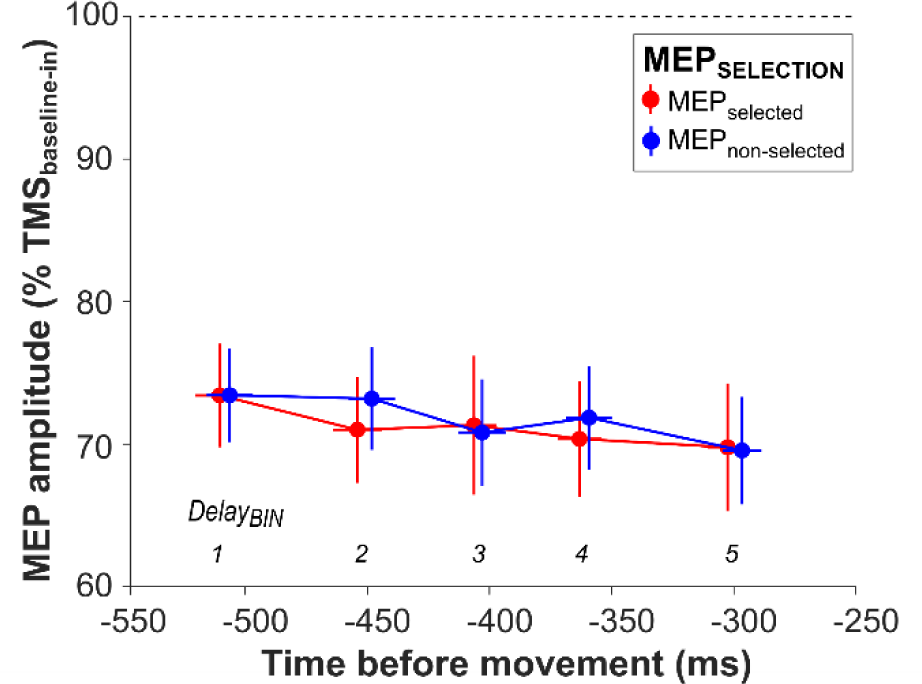
Preparatory suppression according to time before movement onset. MEP amplitudes recorded at TMS_preparation_ (in percentage of MEPs elicited at TMS_baseline-in_) are represented for each Delay_BIN_ in a selected (red) or non-selected (blue) muscle.

### 3.3 Relationship between training-related changes in RTs and CSE

Given that training influenced RTs and CSE, we studied the relationship between changes at these two levels, with CSE considered separately at rest and during action preparation. To assess the relationship between RTs and resting CSE, we ran correlations between training-related changes in RTs and changes in MEPs at TMS_baseline-in_ and TMS_baseline-out_. These analyses did not reveal any link between variations in resting measures of CSE and changes observed in RTs, neither at Training_ratio-early_ **(Fig.7A**, R=-0.27, p=0.36 and R=0.079, p=0.79 for TMS_baseline-in_ and TMS_baseline-out_, respectively) nor at Training_ratio-late_ (R=-0.28, p=0.33 and R=-0.16, p=0.59).

**Figure 7.**
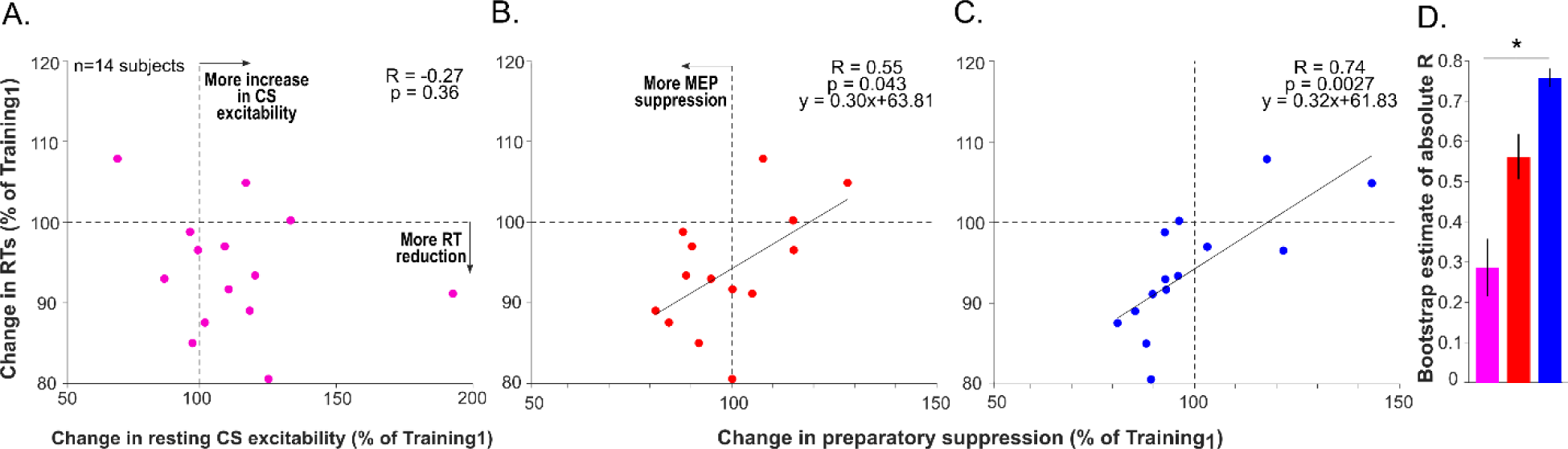
Correlation between early training-related changes in RTs and CSE. Changes in RTs as a function of changes in MEP amplitudes at TMS_baseline-in_ (reflecting resting CSE, **A**) and changes in percentage MEPs at TMS_preparation_ (reflecting preparatory suppression of CSE) in the selected **(B)** and non-selected FDI muscle **(C)** during the early Training_stage_. For this analysis, changes in RTs and MEPs were assessed by computing percentage ratios between the values obtained at Training_3_ and Training_1_. **(D)** Bootstrap estimates of absolute R values are also displayed (± standard deviation of the samples) for each condition. These R values were compared by means of a Pearson and Fillon’s z test. One tail p-values were used given our a priori hypothesis concerning the directionality of the effect (p<0.05).

In contrast, changes in RTs at Training_ratio-early_ were linked to variations in preparatory suppression observed in the selected **(Fig.7B;** R=0.55, p=0.043) and non-selected FDI **(Fig.7C;** R=0.74, p=0.0027): subjects showing a greater training-related strengthening of preparatory suppression also showed larger improvements in RTs. This correlation was not significant at Training_ratio-late_, neither for the selected (R=0.12, p=0.67) nor for the non-selected effectors (R=0.48, p=0.084). Our results suggest that RT improvements were related to early changes in preparatory suppression.

This conclusion is further supported by an additional analysis showing that the strength of the correlation between RTs and CSE at Training_ratio-early_ was significantly higher when considering percentage MEPs at TMS_preparation_ (*i.e.*, preparatory suppression) in the non-selected FDI (bootstrap estimate of absolute R=0.76), than when MEPs were considered at TMS_baseline-in_ (R=0.29; z=1.75; p=0.040, **Fig.7D**). This difference was not significant when taking preparatory suppression in the selected FDI (R=0.56; z-score=0.85, p=0.20). Hence, training-related changes in preparatory suppression of the non-selected effector turned out to be the best predictor of RT improvements.

### 3.4 Single-trial relationship between RTs and preparatory suppression

Finally, we asked whether the dependency of RTs to preparatory suppression is also evident on a single-trial basis. This was the case for MEPs recorded from the non-selected hand: the greater the preparatory suppression in that hand, the shorter the following RT (**Fig.8**, right panel), as supported by the ANOVA_RM_ revealing an effect of the factor MEP_BIN_ on RTs (F_(5,65)_=2.57, p=0.035). Post-hoc tests revealed that RTs in MEP_BIN-1_ and MEP_BIN-2_ (*i.e.*, strongest preparatory suppression) were systematically shorter than those in MEP_BIN-6_ (p=0.0021 and p=0.0090). We did not observe any relationship between RTs and MEPs obtained in the selected hand (MEP_BIN_: GG-corrected F_(2.22,28.88)_=0.85, p=0.45; **Fig.8**, left panel). Hence, the training-related effects and the single-trial relationship indicates that preparatory suppression in the non-selected (non-responding) hand is a predictor of the following RT. The lower this activity, the faster the response.

**Figure 8.**
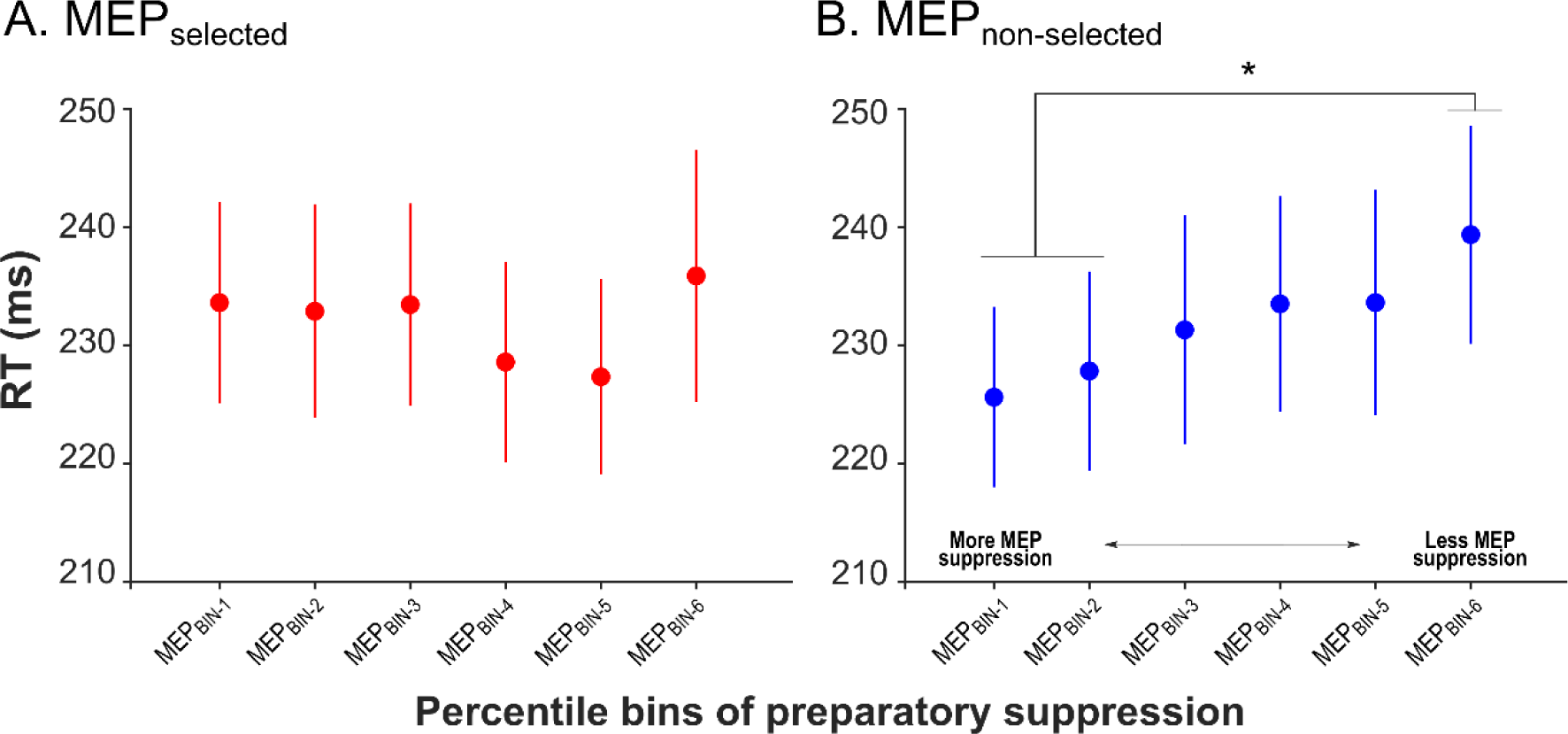
Single-trial MEP-RT relationship. Averaged RTs as a function of the preceding preparatory suppression in a selected (left panel) or non-selected muscle (right panel). For this analysis, the MEP data were divided in 6 MEP_BIN_ of increasing amplitude and the RTs corresponding to each MEP_BIN_ were averaged. The star denotes a significant difference between RTs at MEP_BIN-1_ and MEP_BIN-2_ and RTs at MEP_BIN-6_ in the non-selected muscle (p<0.05). Note that there was also a trend for RTs at MEP_BIN-1_ to be shorter than those in MEP_BIN-4_ (p=0.070) and MEP_BIN-5_ (p=0.066).

## 4. Discussion

Training accelerated RTs while errors remained low. CSE became larger at rest and preparatory suppression of CSE was stronger after training. Interestingly, subjects who showed the strongest RT improvements at the early Training_STAGES_ were also those displaying the largest initial strengthening in preparatory suppression, especially when probed in the non-selected hand. Such a relationship between RTs and preparatory suppression was also evident at a single-trial level: RTs were generally faster in trials where preparatory suppression was deeper.

Subjects responded faster with training. RTs reflect the sum of the time required for processing the imperative cue, preparing the motor command and initiating the action (Derosiere et al. 2019; Haith et al. 2016) and, theoretically, training may impact any of these sensory-motor components. Previous studies have shown that RT improvements can result from both faster sensory processing (Clark et al. 2015) and more efficient motor preparation (Mawase et al. 2018). Yet, in an instructed-delay task, the time required for sensory processing and motor preparation is strongly constrained and most of the RT is assumed to reflect the time needed for action initiation (Haith et al. 2016). Hence, the RT gains reported here are likely to reflect a reduction in initiation time. Our findings thus yield an extension of former work, suggesting that, in addition to accelerating sensory processing and motor preparation, training can also boost action initiation.

Resting CSE was higher when assessed in the context of the task (*i.e.*, at TMS_baseline-in_) than between the blocks (*i.e.*, at TMS_baseline-out_), consistent with previous data (Labruna et al. 2011; Vassiliadis et al. 2018) and with the observation that task-driven increases in attention amplifies cortical excitability (Kastner et al. 1998, 1999). As expected based on prior observations (*e.g.*, (Butefisch et al. 2000; Christiansen et al. 2018; Duque et al. 2008; Galea and Celnik 2009; Pascual-Leone et al. 1995), practicing the task led to an increase in resting CSE. Interestingly, this increase was not exclusive to the task and was in fact strongly similar at TMS_baseline-in_ and TMS_baseline-out_, ruling out the possibility that it resulted from a change in task-related attention over practice (Derosière et al. 2015). Rather, our findings support the idea of a plastic reorganization of the motor system, measurable when engaged in the task as well as at rest.

CSE was reduced during action preparation when compared to baseline (during the task), reflecting the well-known preparatory suppression effect (Duque et al. 2017), which was evident in the selected and non-selected hands from the beginning of the training. Contrary to rest, the amplitude of MEPs at TMS_preparation_ did not increase with practice (they remained unchanged), reflecting a strengthening drop in CSE from the rising baseline state. Notably, although at the group level this reinforcement of preparatory suppression appeared late (**Fig.5A**), at the individual level, a majority of subjects already exhibited a strengthening of preparatory suppression at early training stages (**Fig.6**).

Based on these findings, one could propose that changes in resting excitability are key to RT improvements, as suggested by the inverse relationship between baseline CSE and RTs described recently (Greenhouse et al. 2017). Yet, we did not find a relationship between training-related changes in baseline excitability and improvements in performance. This is in line with the idea that increased resting CSE is not crucial for immediate performance (Bologna et al. 2015), but may be involved in the long-term retention of the motor behavior (Cantarero et al. 2013). Rather, what was predictive of RT gains in the present study was the change in relative CSE, as measured during action preparation: subjects showing the strongest reinforcement of preparatory suppression at the early Training_STAGES_ were those who became fastest. These results are consistent with animal studies showing that behavioral improvements in motor learning tasks are associated with changes in relative preparatory activity (Mandelblat-Cerf et al. 2009; Paz et al. 2003; Vyas et al. 2018). Similarly, a recent study using paired-pulse TMS showed that changes in preparatory activity of M1 intra-cortical circuits are correlated to training-related behavioral gains, contrary to changes observed at rest (Dupont-Hadwen et al. 2018). More generally, our findings agree with the idea that efficient action preparation relies on dynamical shifts of neural activity from a baseline state to a preparatory state (Churchland et al. 2012). From this point of view, training may allow tuning the dynamics of preparatory activity, bringing it closer to an optimal state for action initiation (Vyas et al. 2018). In this line, strengthening of preparatory suppression would facilitate action initiation by allowing excitatory inputs targeting the selected motor representation to better stand out against a quiescent background (mostly reflected in the excitability of non-selected effector), ultimately speeding up RTs (Greenhouse et al. 2015; Hasbroucq et al. 1997; Hasegawa et al. 2017).

This interpretation was reinforced by our single-trial analysis showing that RTs depended on the foregoing amount of preparatory suppression. That is, stronger levels of suppression were related to faster initiation times in the very same trials, in agreement with previous results (Hannah et al. 2018; Hasegawa et al. 2017). Interestingly, we found such relationship when considering the non-selected prime-mover but not the selected one. This was also the case for training-related effects, with preparatory suppression in the non-selected effector appearing as the best predictor of RT changes, possibly because the selected effector is targeted by too many overlapping inputs to supply as meaningful MEP amplitudes (Duque and Ivry 2009). Overall, our data support the view that preparatory suppression facilitates rapid motor initiation.

## Conclusion

This study shows that a simple training paradigm can lead to improvements in action initiation that are accompanied by an increase in resting CSE and a strengthening of corticospinal suppression from the rising baseline state. Moreover, contrary to changes in resting CSE, such strengthening of preparatory suppression was linked to RTs improvements. These findings could have implications for the rehabilitation of patients suffering from impaired action initiation such as in cerebellar ataxia (Battaglia et al. 2006) or Parkinson’s disease (Mure et al. 2012).

## Additional information

### Data availability

The data that support the findings of this study are available at: https://osf.io/8p5wm/ (Vassiliadis 2020).

### Author contributions

PV, GD, JG, JD: conception and design of the work; PV, JG: acquisition of data; PV: analysis of data; PV, GD, JD interpretation of data; PV: drafting; PV, GD, JG, JD revising the manuscript.

